# Functional traits trade-offs define plant population stability worldwide

**DOI:** 10.1101/2022.06.24.497476

**Authors:** Luisa Conti, Enrique Valencia, Thomas Galland, Lars Götzenberger, Jan Lepš, Anna E-Vojtkó, Carlos P. Carmona, Maria Májeková, Jiří Danihelka, Jürgen Dengler, David J. Eldridge, Marc Estiarte, Ricardo García-González, Eric Garnier, Daniel Gómez, Věra Hadincová, Susan P. Harrison, Tomáš Herben, Ricardo Ibáñez, Anke Jentsch, Norbert Juergens, Miklós Kertész, Katja Klumpp, František Krahulec, Frédérique Louault, Rob H. Marrs, Gábor Ónodi, Robin J. Pakeman, Meelis Pärtel, Begoña Peco, Josep Peñuelas, Marta Rueda, Wolfgang Schmidt, Ute Schmiedel, Martin Schuetz, Hana Skalova, Petr Šmilauer, Marie Šmilauerová, Christian Smit, MingHua Song, Martin Stock, James Val, Vigdis Vandvik, David Ward, Karsten Wesche, Susan K. Wiser, Ben A. Woodcock, Truman P. Young, Fei-Hai Yu, Martin Zobel, Francesco de Bello

**Author notes:** **Correspondence: Luisa Conti**, Faculty of Environmental Sciences, Czech University of Life Sciences Prague, Kamýcká 129, 165 00 Praha-Suchdol.

## Abstract

1. Ecological theory posits that temporal stability patterns in plant populations are associated with differences in species’ ecological strategies. However, empirical evidence is lacking about which traits, or trade-offs, underlie species stability, specially across different ecosystems.
2. To address this, we compiled a global collection of long-term permanent vegetation records (>7000 plots from 78 datasets) from a wide range of habitats and combined this with existing trait databases. We tested whether the observed inter-annual variability in species abundance (coefficient of variation) was related to multiple individual traits and multivariate axes of trait variations (PCoA axes).
3. We found that species with greater leaf dry matter content and seed mass were consistently more stable over time (lower variability in species abundance) although other leaf traits played a significant role as well, albeit weaker. Using multivariate axes did not improve predictions by specific traits.
4. Our results confirm existing theory, providing compelling empirical evidence on the importance of specific traits, which point at ecological trade-offs in different resource use and dispersal strategies, on the stability of plant populations worldwide.

## Introduction

Identifying the drivers of temporal stability in plant populations and communities has consequences for maintenance of multiple ecosystem functions over time, including carbon sequestration, fodder resources for livestock, and nutrient cycling (Tilman & Downing, 1994; Hautier *et al*., 2015; Isbell *et al*., 2018). One of the main determinants of community stability is the cumulative temporal variability in the abundances of individual species’ populations (Thibaut & Connolly, 2013; Hallett e*t al*., 2014; Májeková *et al*., 2014). Lower temporal variability in individual population abundances at a given site generally increases overall community stability (Lepš *et al*., 1982, 2018; Pimm, 1984; McCann, 2000). Accordingly, assessing the drivers of temporal variability in populations is necessary to understand and forecast the potential consequences of increasingly common environmental perturbations (Easterling *et al*., 2000; Lloret *et al*., 2012).

While empirical evidence is still scarce and ambiguous, theoretical predictions suggest that the drivers of temporal variability in plant populations are related to different ecological characteristics of species (e.g., r/K life history strategies, MacArthur & Wilson, 1967). These differences can be described through functional traits that determine how plants respond to environmental factors, affect other trophic levels, and influence ecosystem properties (Lavorel & Garnier, 2002; Kattge *et al*., 2011; Garnier *et al*., 2016). Specifically, differences in functional traits among species result in varied responses to the environment that might lead to different patterns of demography, adaptation, and distribution, thus giving rise to different population fluctuations over time (e.g. Angert *et al*., 2009; Metz *et al*., 2010; Adler *et al*., 2013; Májeková *et al*., 2014).

Assessing differences in functional traits between species, as well as the relationship of these differences to specific ecological patterns, has been a long-standing focus in plant ecology leading to a search for general trait trade-offs across taxa and ecosystems (e.g. Díaz *et al*., 2016). Trait trade-offs are generally understood as a shift in the balance of resource allocation to maximise fitness within the constraints of finite resources (e.g. Grime’s C-S-R strategy scheme; Grime, 1977). Traits linked to specific axes of ecological differentiation are key in understanding major trade-offs in plant strategies, such as the trade-off between leaf maximum photosynthetic rate and leaf longevity, also known as the leaf economic spectrum (Wright *et al*., 2004).

Indeed, different specific trade-offs can underlie differences in species’ temporal patterns, both within and between community types. For example, species that are able to respond quickly to environmental variability, e.g. acquisitive resource-use strategy, fast-growing species that invest in organs for rapid resource acquisition and/or high dispersal ability, should sustain higher temporal variation in population size, and will be favoured in sites where disturbance and/or environmental instability determine a fluctuation in resources (MacArthur & Wilson, 1967; Westoby, 1998). In contrast, species adapted to endure environmental variability, e.g. conservative resource-use strategy, slow-growing and long-lived species that invest in structural tissues and permanence, are thought to persist during unfavourable periods due to resources stored from previous, more favourable years (Reich, 2014), and will exhibit less temporal variability (MacArthur & Wilson, 1967; Grime, 2001; Wright *et al*., 2004). These species are expected to be favoured in more stable and predictable environments (Kraft *et al*., 2014).

It remains unclear though whether the potential relationship between species’ traits and species’ stability would be detected through differences in single traits or combined axes of differentiation that incorporate multiple traits (Westoby, 1998; Laughlin, 2014; Díaz, *et al*. 2016). Several ecological strategy schemes, such as the classic r/K selection (MacArthur & Wilson, 1967) and C-S-R (Grime, 1977) theories, as well as the Leaf-Height-Seed scheme (’
sLHS’; Westoby, 1998), can theoretically help predicting how functional trade-offs determine species’ temporal strategies and their fitness across different types of environments. The LHS scheme for instance, is based on three independent plant traits which should provide key proxies for independent trade-offs in plants (stress adaptation, competition, and response to disturbance respectively; Westoby, 1998). Interestingly, only a few empirical studies have linked differences in temporal strategies to functional traits within plant communities (Adler *et al*., 2006; Angert *et al*., 2009; Metz *et al*., 2010; Májeková *et al*. 2014; Craven *et al*., 2018). For example, Májeková *et al*. (2014) empirically confirmed that herbaceous species with a more conservative resource-use strategy (i.e., those with higher leaf dry matter content - LDMC) have more stable populations over time. A similar relationship was found at the community level, where communities including a greater abundance of species with high LDMC were more stable (Polley *et al*., 2013; Chollet *et al*., 2014). A recent global meta-analysis of sown grasslands suggested that an increase in the abundance of rapidly-growing species can destabilize community biomass over time (Craven *et al*., 2018). This is supported by empirical demonstrations that community stability is predicted by the functional traits of the dominant species rather than by species diversity *per se* (Lepš *et al*., 1982). Further, only Májeková *et al*. (2014) tested whether trait-based predictions of population temporal variability were consistent across different managements, i.e. fertilization and competitor-removal treatments, generally finding minor differences and consistent predictions for LDMC. Ultimately, global empirical evidence of a general link between quantitative functional traits and the temporal variability of populations, and whether this link is maintained despite differences in community types and environmental conditions, is still missing (de Bello *et al*., 2021).

Here, using a global compilation of long-term, recurrently monitored vegetation plots, encompassing different habitat types (https://lotvs.csic.es/), we determine which plant traits better predict the temporal stability of plant populations. We expect that populations of species with more acquisitive and higher dispersal-ability traits will tend to be more variable over time, while those of species with more conservative trait values and lower dispersal ability will tend to be more stable over time. We also expect to find global empirical evidence of the generality of these relationships.

## Materials and Methods

### Plots and population’s stability

We used 78 datasets consisting of a total of 7396 permanent plots of natural and semi-natural vegetation that have been consistently sampled for periods of between six and 99 years, depending on the dataset (Supporting Information Table **S2**; Valencia *et al*. 2020a, Sperandii *et al*. 2021). These datasets were collected from study sites around the globe, and differ in sampling method (e.g., above-ground biomass, visual species cover estimates, species individual frequencies), plot size, and study duration. The studies that generated the datasets sampled different types of vegetation and covered a wide array of biomes, with mean annual precipitation spanning from 140 mm to 2211 mm, highest temperature of the warmest month spanning from 11.3°C to 35.7°C, and lowest temperature of the coldest month spanning from -35.3°C to 7.7°C (Supporting Information Table **S2**).

First, for each plot we quantified the inter-annual variability in the size of each species’ population using the coefficient of variation (CV) of abundance over time, i.e. the standard deviation of species abundance over mean species abundance (Májeková *et al*., 2014; de Bello *et al*., 2021). Since a fundamental differentiation between growing strategies corresponds to woody versus non-woody species (Reich, 2014; Díaz *et al*., 2016), here we focused on non-woody species, thus excluding forest overstories and woody species’ seedlings when present. Moreover, based on the collected data available, we could not distinguish adult woody individuals from seedlings, with seedlings most likely being the cause of high variability in woody species’ CV values (see Supporting Information Fig. **S1a**). To avoid using biased CV values (increased CV for very sporadic species), we excluded those species that occurred in fewer than 30% of the sampling events across the time series for a given plot (Májeková *et al*., 2014). Further, to account for variability in CV values between and inside the datasets, mostly due to differences in abiotic, biotic, and management conditions, we calculated the average CV value for each species in each dataset, standardizing and scaling these averages within each dataset (z-scores). This resulted in a total of 3,397 species *per* dataset CV values. To account for potential effects of temporal directional trends in vegetation affecting CV (Valencia *et al*., 2020b) we also computed a detrended version of CV (CVt3) which gave very similar results to the basic CV calculations (see Supporting Information Fig. **S2**).

### Functional traits

For all the species in our dataset, we collected trait information from the TRY global database (Kattge *et al*., 2020). We specifically focussed on continuous traits of herbaceous species, which were more appropriate for depicting trait trade-offs along axes of species differentiation (see Supporting Information Fig. **S1**). Specifically, we analysed plant height, seed mass, specific stem density, LDMC, specific leaf area (SLA), leaf nitrogen content *per* unit mass, and leaf phosphorus content *per* unit mass (see Garnier *et al*., 2017 for trait name nomenclature and definitions). For each species, we averaged trait values across all standard measurements obtained from TRY, excluding those performed under explicit treatments, on juveniles, and outliers. The traits were log-transformed when their distribution was skewed (using natural logarithm). For details on the traits used, their summary statistics, their correlations, and their coverage in each dataset, see Supporting Information Table **S3**. To take into account multivariate trade-offs between species, we also considered axes of functional variation derived from multivariate analyses (Principal Coordinates Analysis, PCoA). The traits considered were weakly inter-correlated, with the two major axes of trait differentiation from PCoA, linked mainly to LDMC and seed mass (see Supporting Information Table **S1** for details). The taxonomic names follow the nomenclature of ‘The Plant List’ (www.theplantlist.org). Nomenclature was standardized using the R package ‘Taxonstand’ (Cayuela *et al*., 2017).

### Data analyses

To quantify how the considered continuous traits were linked to species CV, we used linear mixed effect models (‘lmer’ function in R package “lme4”, Bates *et al*., 2014). As a response variable, we used the mean CV for each species in each dataset, standardized as mentioned above. As predictors, we included all the continuous traits listed above, scaled and centered. To account for the taxonomic and spatial structure of the data, we included both species identity and dataset identifier as random intercept factors. We visually checked the compliance of model residuals with normality and homoscedasticity. To assess the goodness-of-fit of the full model, marginal and conditional R^2^ were calculated (Nakagawa & Schielzeth, 2013; Nakagawa *et al*., 2017). Then, we compared the marginal R^2^ of nested models, each differing in the subset of predictors that were included; therefore, we fitted a final model that included a subset of the original predictors used and had the highest marginal R^2^. Similar models were run using, instead of single traits, both the multivariate PCoA axes that resulted from the combination of traits. We also fitted separate models using each a single trait of those emerging as stronger in the final multivariate model (See Supporting Information Table **S1**). In addition, to explore the consistency of the stability-trait relationships across datasets, we fitted models using each single trait and adding random slope effect for the datasets (Supporting Information Fig. **S4**). Finally, similar models were run also on the two components determining species’ CV separately, i.e. mean abundance and standard deviation of abundance in time, also standardizing these variables within each dataset (Supporting Information Fig. **S3**).

## Results

We were able to detect two sets of key continuous functional traits playing a consistent major role in species’ population temporal stability: one linked exclusively to seed mass, and the other linked to the leaf economic spectrum, i.e. LDMC, SLA, and Leaf N content. Based on the final linear mixed effect model, these two sets of traits had the most influence on species CV among the continuous traits considered (Table **1**; Figure **1**, Figure **2**).

**Table 1.** Model’s summary for both the full model, containing all the predictors, and the final model, containing only a subset of the initial predictors. Estimates and relative standard errors are shown. R^2^ (fixed): variation explained by fixed factors; R^2^ (total): variation explained by both fixed and random factors. P-values calculated using Satterthwaite approximation for degrees of freedom. ***p-value<=0.001; **p-value<=0.01; *p-value<=0.05.

|  | Full model | Final model |
| --- | --- | --- |
| (Intercept) | -0.10<br>(0.06) | -0.03<br>(0.04) |
| Plant height | -0.01<br>(0.09) |  |
| Leaf N content | 0.03<br>(0.08) | 0.06<br>(0.04) |
| Leaf P content | 0.04<br>(0.07) |  |
| Seed mass | -0.12<br>(0.08) | -0.08 *<br>(0.04) |
| SLA | 0.02<br>(0.09) | 0.09 *<br>(0.04) |
| LDMC | -0.23 **<br>(0.07) | -0.21 ***<br>(0.04) |
| SSD | 0.06<br>(0.06) |  |
| N | 676 | 1630 |
| Species | 93 | 395 |
| Datasets | 67 | 77 |
| $R^2$ (fixed) | 0.05 | 0.07 |
| $R^2$ (total) | 0.13 | 0.18 |

**Figure 1.**
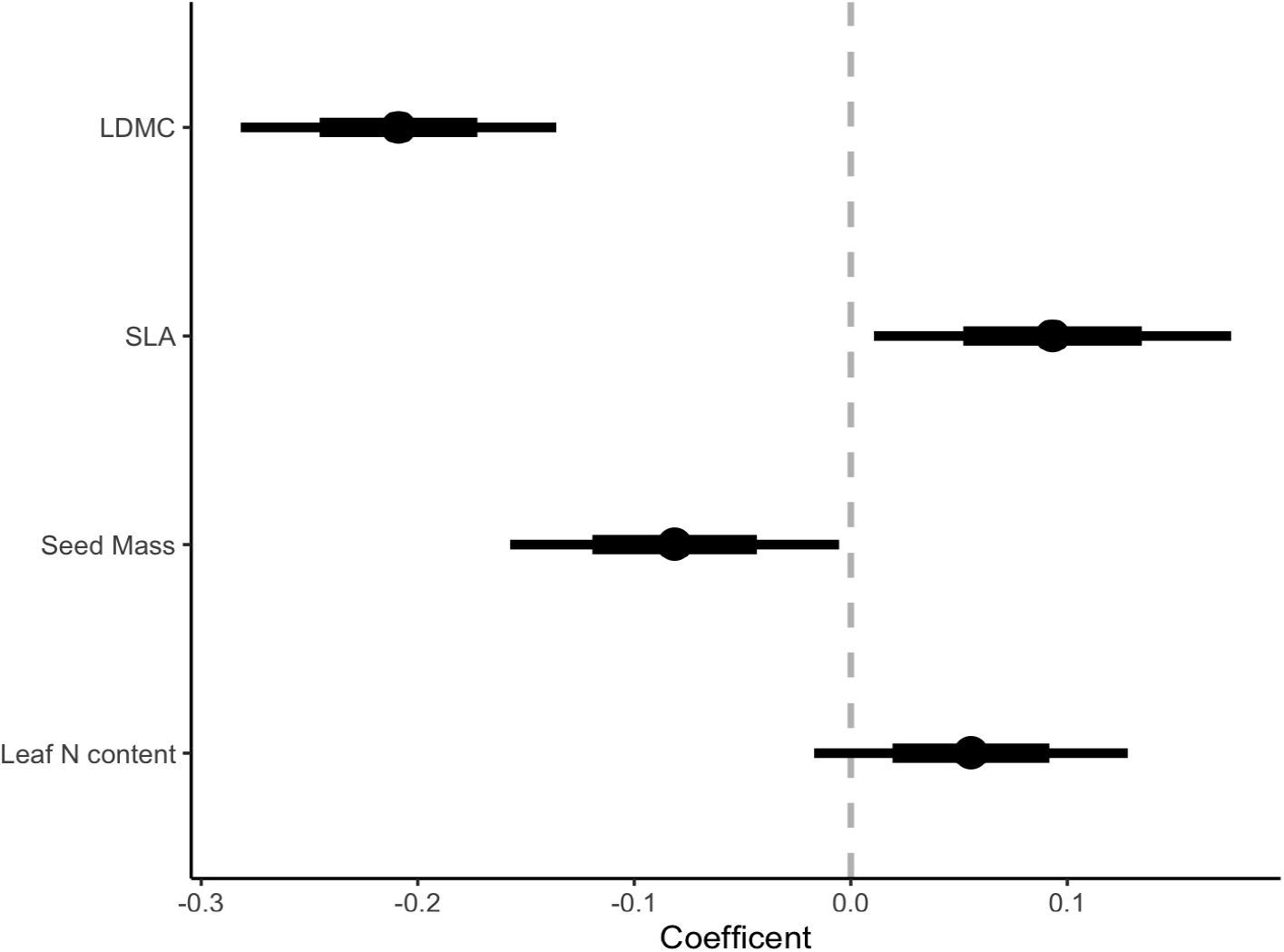
Coefficient plot showing estimate values and their 68% (thin line) and 95% (thick line) confidence intervals of the final linear mixed effect model fitted. To explain species CV, the final model included leaf dry matter content (LDMC); seed mass transformed through natural logarithm (Seed Mass); specific leaf area transformed through natural logarithm (SLA); and Leaf N content.

**Figure 2.**
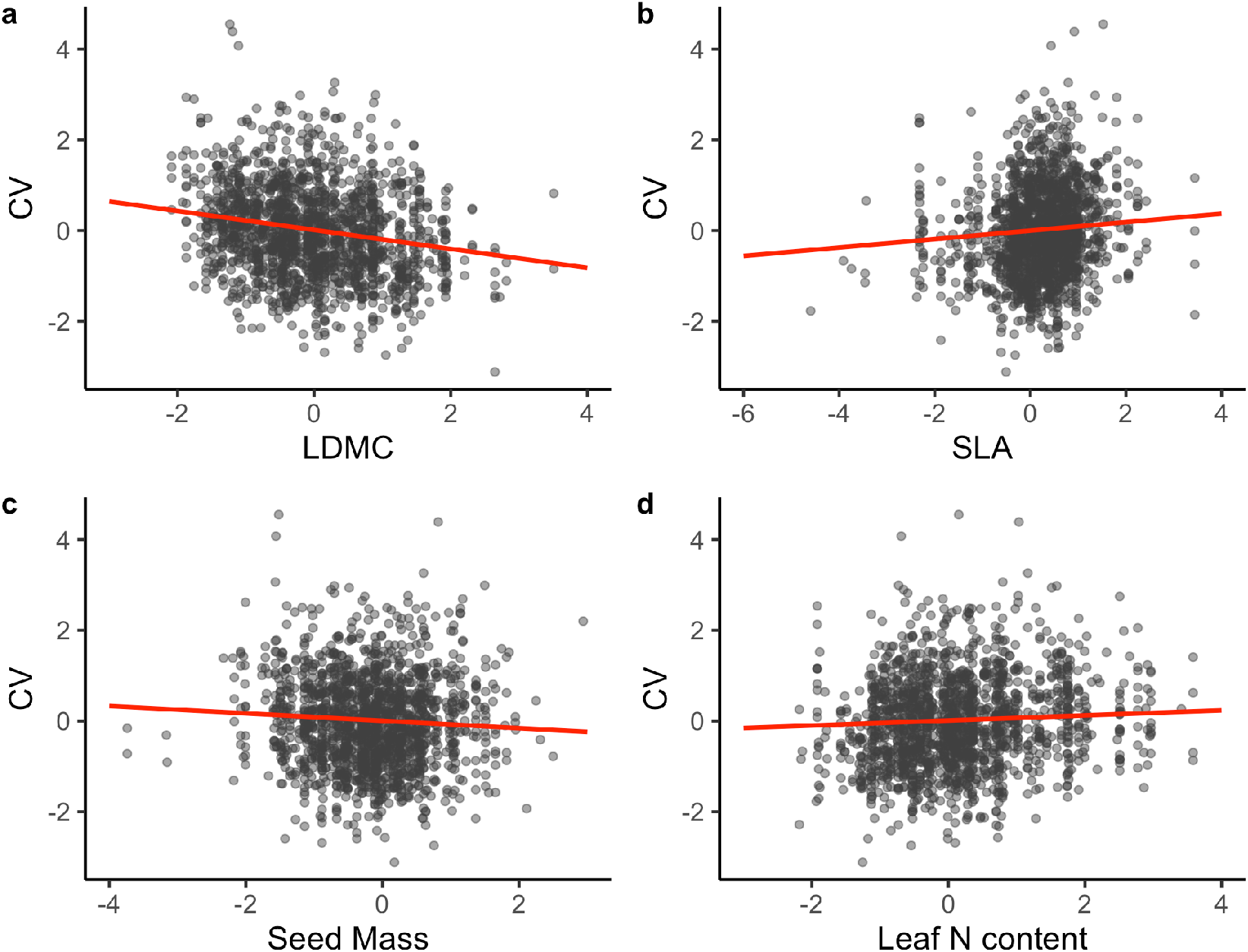
Regression plots of the final model showing the effects of leaf dry matter content (LDMC, a), specific leaf area (SLA, b), seed mass (c), and leaf N (d) content on the CV of species.

We found significant negative coefficients with population variability for LDMC and for seed mass (Table **1**; Fig. **1**). These coefficients indicate that species with greater LDMC and greater seed mass were more stable (i.e. lower CV values; Fig. **2a**). In contrast, we found positive coefficients for SLA and Leaf N content, although the effect was statistically significant only for SLA. For these traits, the larger the trait value, the higher the species CV and therefore the less stable the species populations (Fig. **2b**,**d**). It should be noted that the effect of these traits was rather consistent across datasets (low deviation of the datasets’ random slope effect compared to the main effect slope for both the models using LDMC and seed mass; Supporting Information Fig. **S4**).

Similar results were found using either of the two first PCoA axes based on multiple traits (Supporting Information Table **S1**), although with a slightly lower predictive power (R^2^ fixed was 0.05 compared to 0.07 in the final model with individual traits). Therefore, results from PCoA did not improve results from single traits. We also fitted models using the single PCoA axis and the single traits. In this case as well single trait models explained a higher variability compared to the models with the single PCoA axis (PCoA Axis 1 model’s R^2^ fixed was 0.040 vs 0.050 when using LDMC; PCoA Axis 2 model’s R^2^ fixed was 0.003 vs 0.005 when using seed mass; Supporting Information Table. **S1**). Finally, when the two components determining species’ CV were analysed separately, i.e. species’ mean abundance and standard deviation of abundance over time, the model predicting mean abundance was stronger compared to the one for standard deviation of abundance over time (with significant results and a higher R^2^ fixed; see Supporting Information Fig. **S3**) although LDMC predicted significantly both mean abundance and its standard deviation.

## Discussion

By analysing a large worldwide compilation of permanent vegetation plot records, we confirmed the generality of theoretical predictions relating key functional traits to plant population stability over time. We specifically found that the abundance of species with greater LDMC and a bigger seed mass were the most stable over time. Ultimately, these results suggest that common functional trade-offs related to resource use and dispersal consistently define herbaceous plant population stability worldwide.

We identified two likely functional trade-offs that influence stability. Specifically, differences associated with the leaf economic spectrum (in our case linked to LDMC, SLA and N content values) define trade-offs in terms of slow-fast resource acquisition (Wright *et al*., 2004; Díaz *et al*., 2016). Differences in seed mass values represent the competition-colonization (seedling establishment) trade-off (Turnbull *et al*., 1999) related to the species’ dispersal and establishment strategy. Moreover, when analysing multivariate functional differentiation in herbaceous species, these set of traits were the ones most strongly associated with the two first principal axes (Supporting Information Table **S1**), further confirming the importance of these two functional differentiation axes. These findings are broadly consistent with Diaz *et al*. (2016), who found that the main differentiation between species was related to leaf and size-related (whole plant and seed) traits. At the same time, it is interesting to notice that, in our case, combined trait information in the form of plant spectra (i.e. via the PCoA axes) lost some ecological explanatory power compared to specific trait effects. This suggests that, in the case of predicting species stability, using specific functional traits could be more effective than using axes of functional variation based on multiple traits, in which case their individual effects could be possibly blurred.

Both the two main functional traits ultimately related to the populations’ temporal patterns are intrinsically linked to how the species adapt to patterns of resource availability and disturbance. Higher LDMC values, as well as smaller SLA and N content values, correspond to a slow return of investments in nutrients, lower potential relative growth rate, and longer leaf and whole-plant lifespan (Wright *et al*., 2004; Pérez-Harguindeguy *et al*., 2013). This implies higher potential of buffered population growth. In fact, slow-growing and long-lived species, for example with higher values of LDMC, could have an advantage in unfavourable years due to resources stored from previous, more favourable years, thus maintaining buffered population growth and consequently more stable populations (Májeková *et al*., 2014; Reich, 2014). Similarly, larger seed mass means greater resources stored that tend to help the young seedling establish and survive in the face of stress with the cost of short-distance dispersal, while smaller seeds (also in combination with seed shape) are typically related to greater longevity in seed banks and dispersal over longer distances (Thompson *et al*., 1993; Turnbull *et al*., 1999; Moles & Westoby, 2006). Therefore, species germinating from seeds with a larger mass are more likely to survive during adverse years and so their populations are more stable in a given site compared to species with smaller seeds, which will tend to maintain their populations through permanence in seed banks, which enables proper germination timing (Venable & Brown, 1988; Metz *et al*., 2010). In addition, species with greater seed mass might be favoured in communities where gaps are scarce, which are usually dominated by perennial species (with higher LDMC values) and are more stable. Large seeds will tend to remain closer to the mother plant than small seeds, thus increasing the stabilizing effects on populations. Small seeded species still maintain a buffered population growth (Pake & Venable, 1995), yet their above-ground abundance will be more variable over time, because they usually germinate only in favourable years. This explanation is particularly supported, for example, for short-lived plants (annuals and biennial species together, Table **S3**), which tend to be less stable over time (Fig. **S1b**) and are generally associated with the small-seed strategy at a global scale (Westoby, 1998).

It is important to consider that the same traits that predicted species variability, using CV, also predicted the components of CV, i.e. species means and standard deviation (SD). Clearly the SD in species fluctuation is inherently increasing with species means, following the so-called Taylor’s power law (Lepš, 2004). This leads to the use of CV in the study of stability, as a more “scaled” measure of species variability. At the same time, when the CV is negatively correlated to species means, as in our case (R=-0.46, which corresponds to the case of a slope in the Taylor’s power law being lower than 2), it implies that more dominant species tend to fluctuate comparatively less than subordinate species. This is an important observation because this scenario implies that the same type of species that are dominant, e.g. with high LDMC, are also the more stable ones. Since dominant species were key drivers of the stability of the communities considered in our study (Valencia *et al*., 2020a) the results of the present study indicate that the same traits that determine species dominance also determine species stability, which is a key message for any attempt to predict both community structure and its potential to buffer environmental fluctuations (de Bello *et al*., 2021).

Our results show worldwide evidence that species with more conservative leaf economics and greater seed mass are generally more stable, i.e. less variable over time, and therefore confirm theoretical assumptions as well as previous localized empirical evidence on the interdependence between these traits, their relative trade-offs, and population temporal stability (e.g. MacArthur & Wilson, 1967; Májeková *et al*., 2014). In addition, our results show the global validity of these trade-offs, found across a variety of abiotic and biotic conditions. Overall, our findings contribute to a better understanding of the drivers of plant population temporal stability, which has important implications for the conservation of ecosystem functions over time across the world.

## Supporting information

Table S3 Functional traits information

## Acknowledgements

This research was funded by Czech Science Foundation Grant GACR16-15012S and Czech Academy of Sciences Grant RVO 67985939. RJP was supported by the Scottish Government’s Rural and Environmental Sciences and Analytical Services division. MP and CPC were supported by the Estonian Research Council grant (PRG609, PSG293). MP and MZ were supported by the European Regional Development Fund (Centre of Excellence EcolChange). SKW was supported by the Strategic Science Investment Fund of the New Zealand Ministry of Business, Innovation and Employment. EV was funded by the 2017 program for attracting and retaining talent of Comunidad de Madrid (n° 2017-T2/AMB-5406). RM was supported by Defra and the Leverhulme Trust.

## Author Contributions

FdB and EV conceived the idea together with LC, EV and TG gathered the data, LC prepared the data, performed the analyses, and wrote the first draft of the manuscript. LG, JL, AE-V, CC, and MM, helped with data preparation and/or statistical analyses. The rest of the authors contributed with data. All the authors actively participated in the writing.

## Data Availability

All the metrics used in the analyses are available at https://doi.org/10.5281/zenodo.6720583 under CC-BY licence. For access to the LOTVS datasets in full please refer to https://lotvs.csic.es/

The following Supporting Information is available for this article:

**Fig. S1** Mean species variability (CV) and categorical traits

**Fig. S2** Mean species variability detrended (CVt3) and traits

**Fig. S3** Species’ mean abundance and standard deviation, and traits

**Fig. S4** Random slope effects in single trait models.

**Table. S1** Mean species variability (CV) and PCoA axes

**Table S2** Dataset information (Separate file: “Table S2 Datasets information.xlsx”)

**Table S3** Functional traits information (Separate file: “Table S3 Traits information.xlsx”)

## Supporting Information

**Fig. S1.**
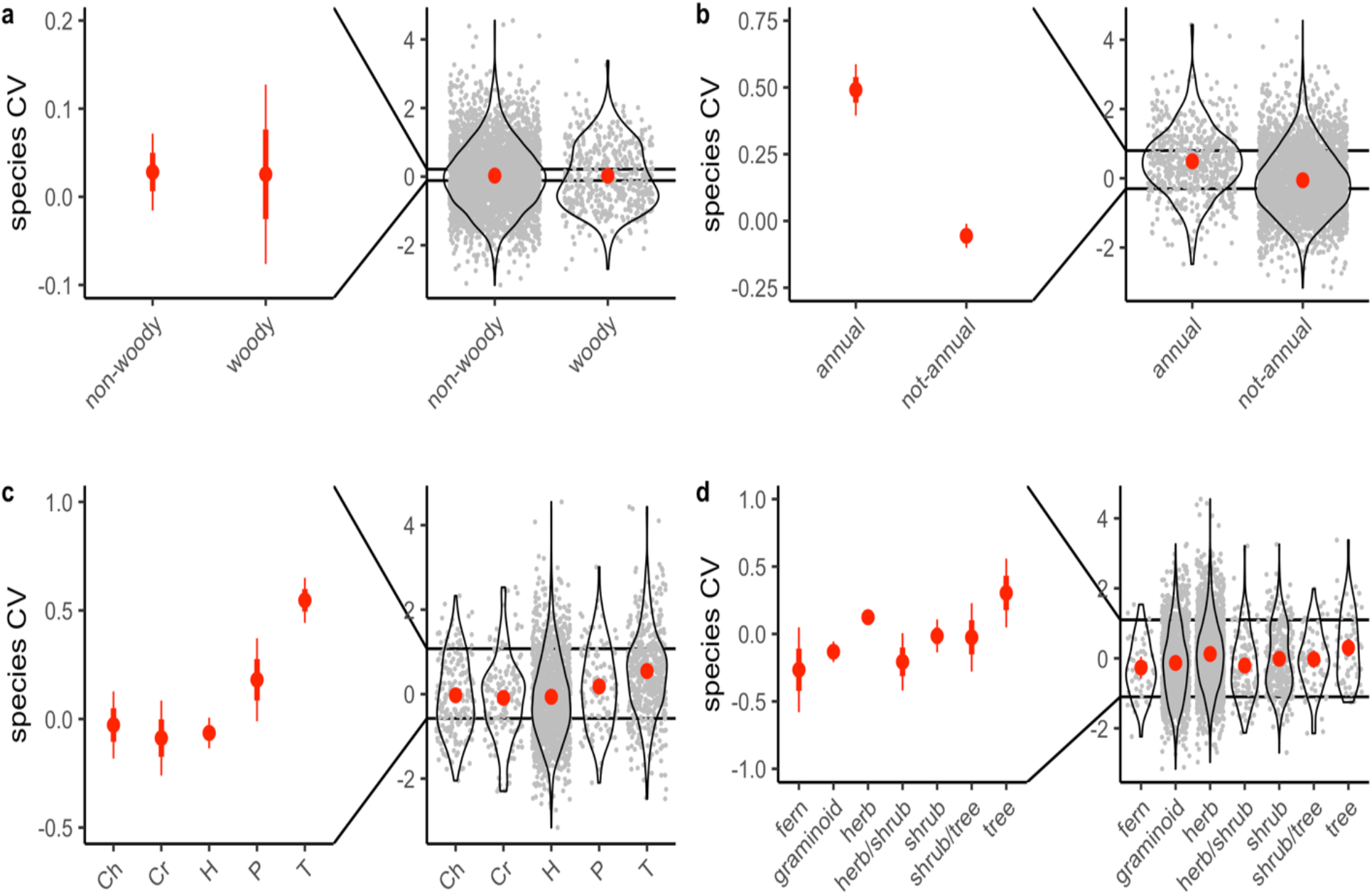
Species variability (CV) and categorical traits. Here we show results of the models fitted using single categorical traits as predictors for the mean species CV at dataset level (i.e. analogous models as the final model in the main text): woodiness (a), life span (b), life form (c), growth form (d). Coefficient plots of these linear mixed models are shown in red (estimates and respective 95% confidence intervals). Intercept was excluded from the model to better understand the differences across trait categories.

**Fig. S2.**
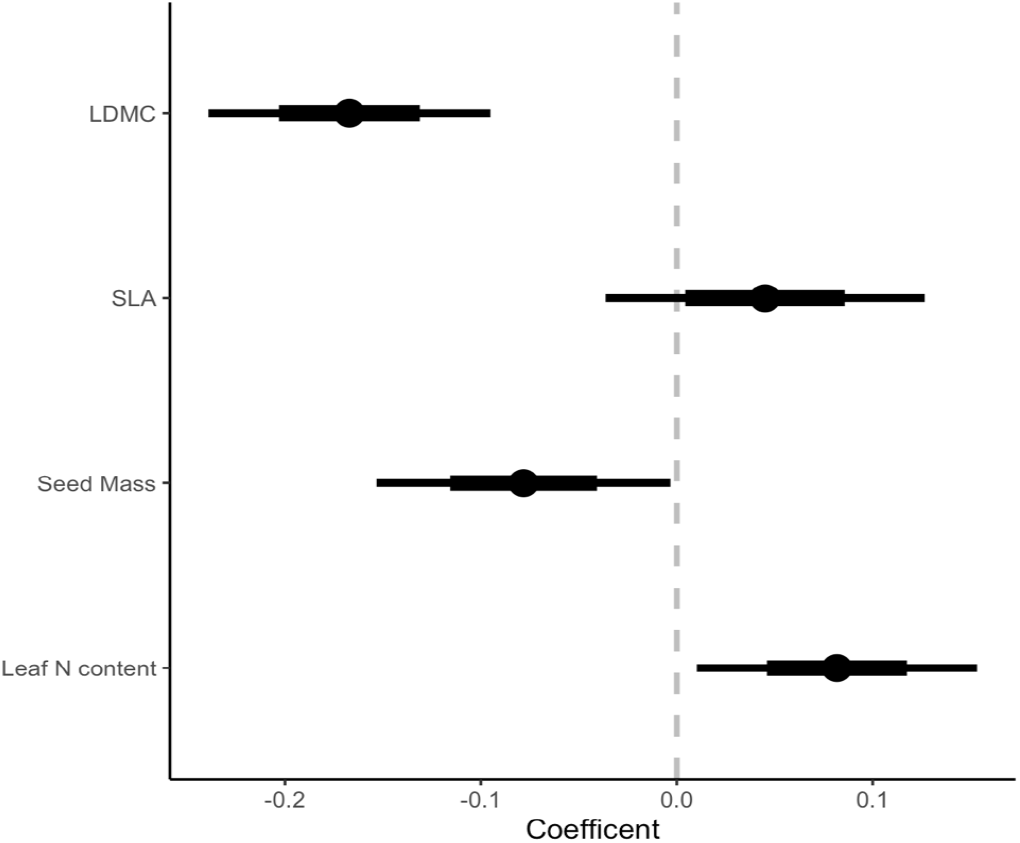
Mean species variability detrended (CVt3) and traits. We computed the detrended version of CV using the moving window method in Valencia *et al*. (2020b). We then fitted a model analogous to the final model in the main text. Results were very similar to those in the main text and are not further discussed. In this model, R^2^ (fixed) was 0.05 while R2 (total) was 0.16.

**Fig. S3.**
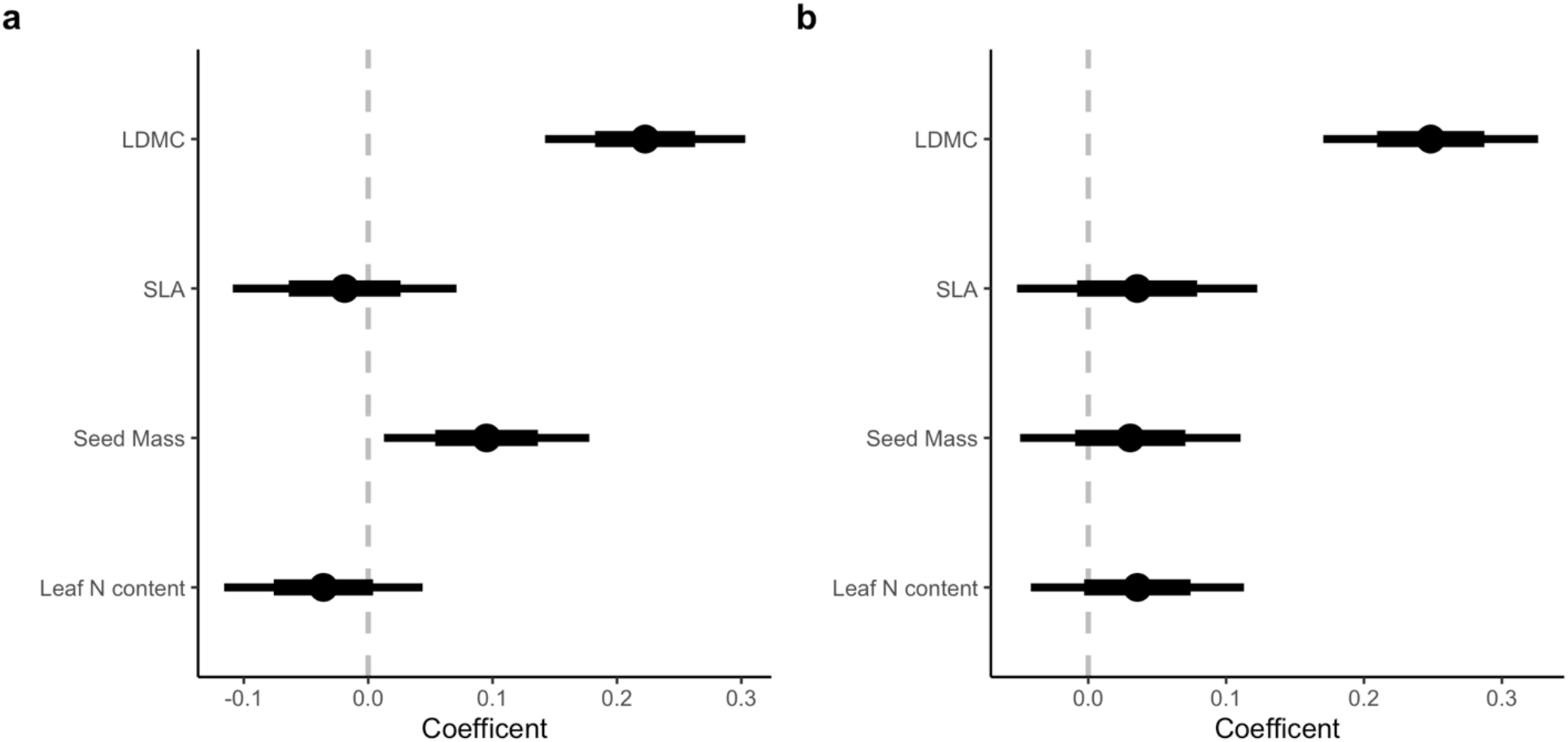
Species’ mean abundance and standard deviation, and traits. We fitted analogous models as the final model in the main text but using either the mean abundance of each species in each dataset (a), or their standard deviation (b), both these variables where scaled and centered within each dataset. The model using the mean abundance was stronger, with more significant results and higher R^2^ (fixed 0.05, total 0.22), compared to the model using the standard deviation (R^2^ fixed 0.04, total 0.19). Moreover, the positive effect of LDMC on the species’ standard deviation is due to the known relationship between variation and mean abundance (Pearson’s correlation coefficient between CV and mean abundance is -0.46).

**Fig. S4.**
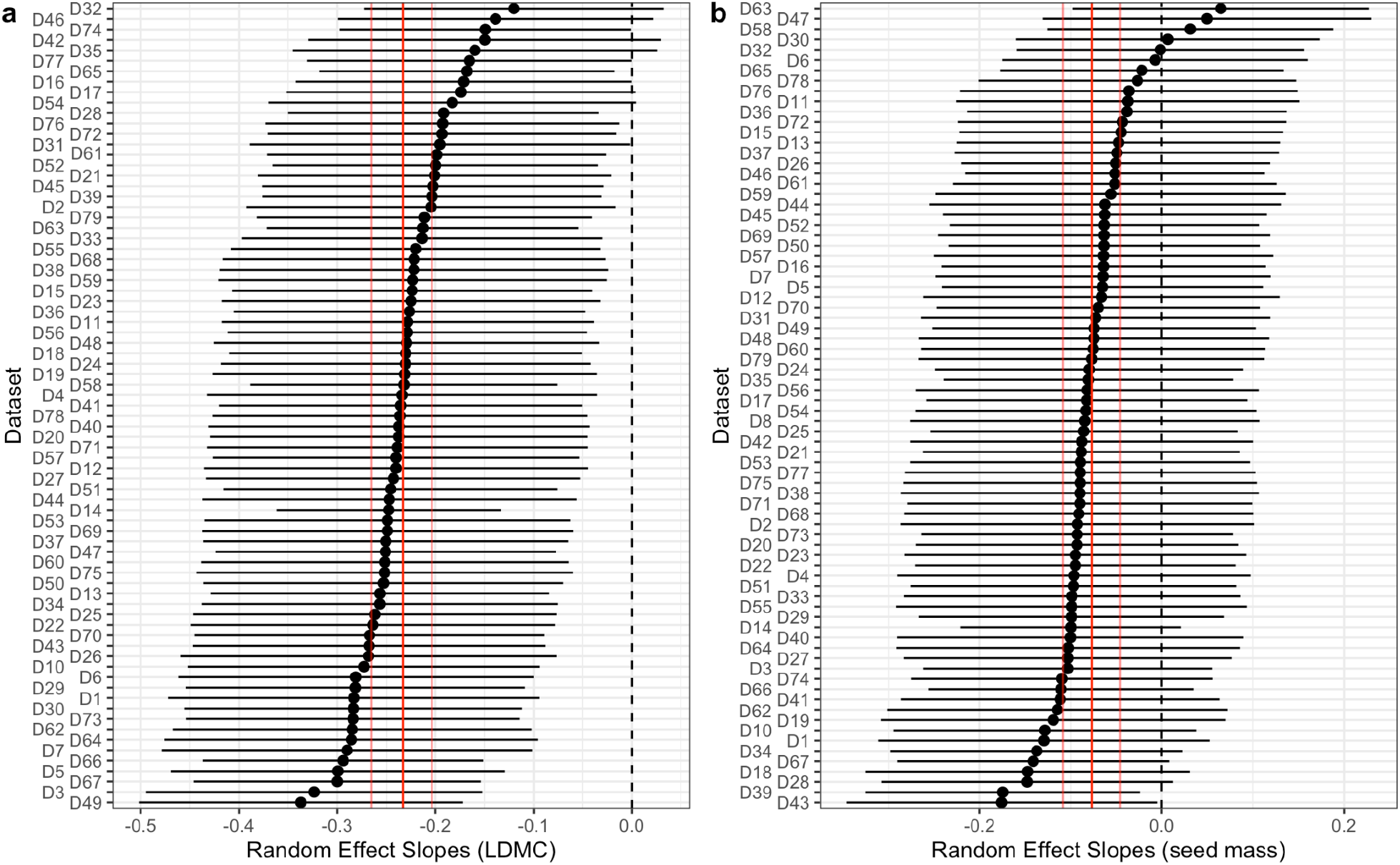
Random slope effects in single trait models. To see the variability across datasets of the relationships found in the main results, we fitted two separate models explaining mean (dataset level) species variability (CV) with each the two main traits emerging from the final model in the main text, i.e leaf dry matter content (LDMC) and seed mass, adding a random slope effect. Here, caterpillar plots show the resulting random effect slope for each of the datasets analyzed in the model using LDMC (a) and the model using seed mass (b).

**Table. S1.**
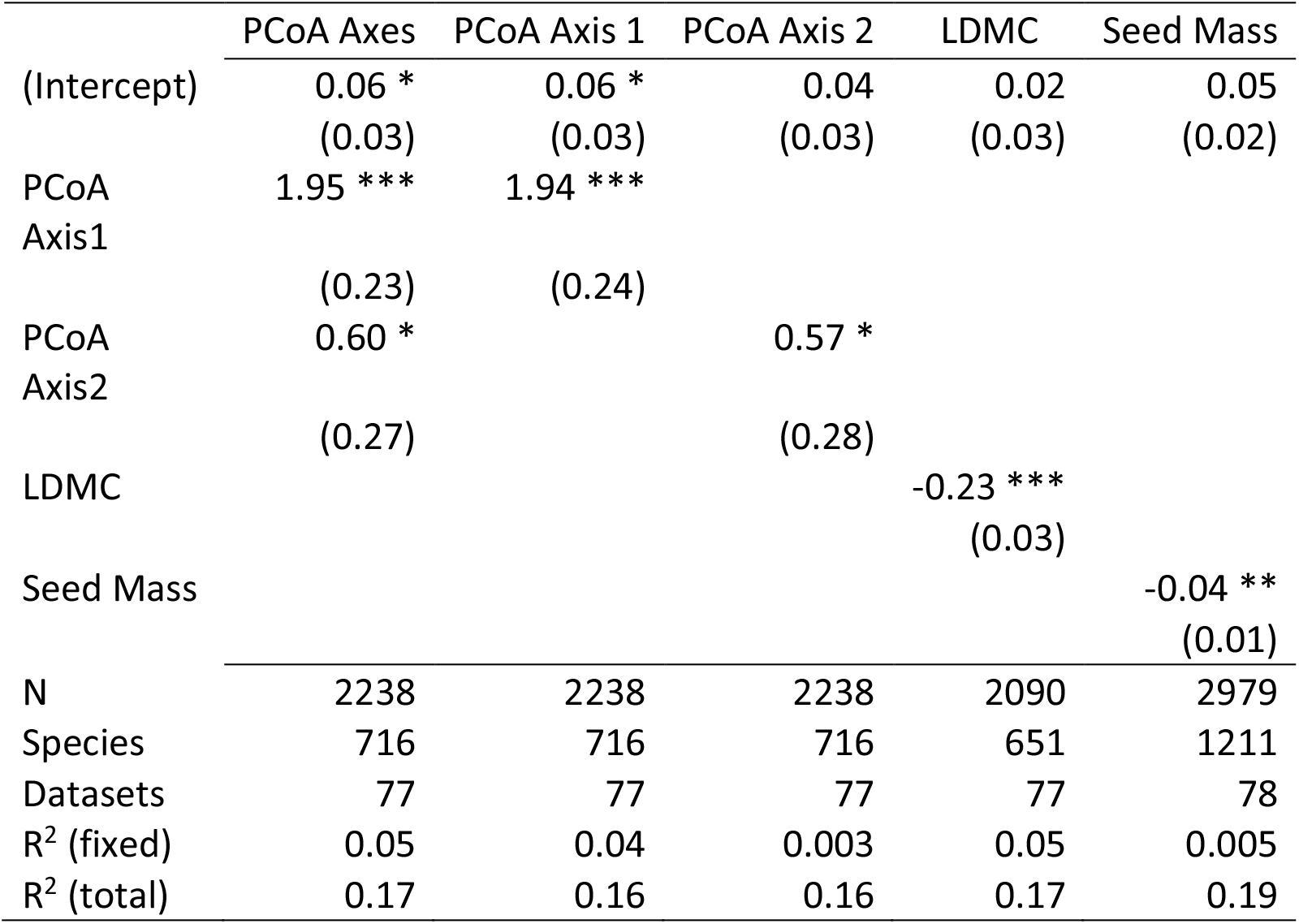
Mean species variability (CV) and PCoA axes and single traits. We performed a PCoA considering all continuous traits. We used a PCoA instead of a PCA as we couldn’t use a correlation-matrix PCA because of the missing trait data. Trait data was centered and scaled, as well as log-transformed when its distribution was skewed. We used Gower’s distance to generate the pairwise distance matrix, which was corrected through squared-root transformation. We found that the first two axes resulting from the PCoA explained 15% and 14% of the variability (when considering only positive eigenvalues, i.e. the metric part of the distance matrix, which explained 82% of the total variability). Moreover, these two axes were highly correlated (r>0.8) to leaf dry matter content (LDMC) and seed mass, respectively. We fitted a linear mixed effect model analogous to the final model in the main manuscript but using the values from these two axes. We found that despite the high correlation with the traits explaining CV in the main results, the results using the PCoA axis were not as clear as when using the trait values. Their R^2^ (fixed) value was of 0.05, compared to 0.07 in the main results (Tab.1). For a fair comparison, we also fitted models using the single PCoA axis and the single traits, also in this case the single trait models explained a higher variability compared to the models with the single PCoA axis. All predictors were mean-centred and scaled by 1 standard deviation, to be able to compare results across all models. R^2^ (fixed): variation explained by fixed factors; R^2^ (total): variation explained by both fixed and random factors. P-values calculated using Satterthwaite approximation for degrees of freedom. ***p-value<=0.001; **p-value<=0.01; *p-value<=0.05.

